# Control of alveolar bone development, homeostasis, and socket healing by salt inducible kinases

**DOI:** 10.1101/2024.09.04.611228

**Authors:** Nicha Tokavanich, Byron Chan, Katelyn Strauss, Christian D. Castro Andrade, Yuki Arai, Mizuki Nagata, Marc Foretz, Daniel J. Brooks, Noriaki Ono, Wanida Ono, Marc N. Wein

## Abstract

Alveolar bone supports and anchors teeth. The parathyroid hormone-related protein (PTHrP) pathway plays a key role in alveolar bone biology. Salt inducible kinases (SIKs) are important downstream regulators of PTH/PTHrP signaling in the appendicular skeleton where SIK inhibition increases bone formation and trabecular bone mass. However, the function of these kinases in alveolar bone remains unknown. Here, we report a critical role for SIK2/SIK3 in alveolar bone development, homeostasis, and socket healing after tooth extraction. Inducible SIK2/SIK3 deletion led to dramatic alveolar bone defects without changes in tooth eruption. Ablating these kinases impairs alveolar bone formation due to disrupted osteoblast maturation, a finding associated with ectopic periostin expression by fibrous cells in regions of absent alveolar bone at steady state and following molar extraction. Distinct phenotypic consequences of SIK2/SIK3 deletion in appendicular versus craniofacial bones prompted us to identify a specific transcriptomic signature in alveolar versus long bone osteoblasts. Thus, SIK2/SIK3 deletion illuminates a key role for these kinases in alveolar bone biology and highlights the emerging concept that different osteoblast subsets utilize unique genetic programs.

**Summary statement:** SIK2/SIK3 deletion in alveolar bone reduces bone formation and mass by impairing osteoblast maturation, unlike in long bones, where it increases bone formation and mass.

## Introduction

Alveolar bone surrounds the teeth in the mandible and maxilla. Together with periodontal tissues, including periodontal ligament, cementum, and gingival tissue, alveolar bone is essential for supporting teeth and facilitating mastication. Like other skeletal tissue, it serves as a mineral reservoir and is produced by mesenchymal-lineage osteoblasts. Unlike other skeletal tissues, alveolar bone originates from neural-crest-derived dental follicle cells and primarily forms through intramembranous ossification^1^. The development and maintenance of alveolar bone are believed to rely on the presence of teeth^2^. Alveolar bone and the periodontium develop from complex interactions between dental epithelium and ectomesenchyme during tooth eruption^3–6^. Injury or degeneration of alveolar bone can lead to tooth loosening or loss, detrimentally affecting quality of life^7^. Due to its complex structure, alveolar bone is challenging to regenerate, making it vital to understand its development, maintenance, and healing in order to treat periodontal disease and regenerate periodontally compromised tissue.

Calciotropic endocrine hormones such as parathyroid hormone (PTH) and calcitonin can influence alveolar bone metabolism^8^. The PTH and parathyroid hormone-related protein (PTHrP) pathways are crucial in tooth and bone development, mineral ion homeostasis, and bone remodeling^5,9–15^. During craniofacial development, paracrine PTHrP signaling regulates tooth eruption and tooth root development. PTHrP-expressing dental follicle cells differentiate into cementoblasts, periodontal ligament cells, and alveolar bone osteoblasts^9,10^. Autocrine signaling through the PTH/PTHrP receptor (PTH1R) ensures that dental follicle mesenchymal cells develop into functional periodontium, facilitating normal tooth eruption. PTH1R deficiency causes an abnormal shift in the fate of PTHrP-positive mesenchymal progenitor cells, resulting in the formation of cementoblast-like cells rather than cells of the functional periodontal apparatus, impairing tooth root formation, periodontal ligament development, and tooth eruption^10^. Loss-of-function PTH1R mutations in humans causes Primary Failure of Eruption (PFE), an autosomal dominant disorder characterized by halted tooth eruption despite an unobstructed eruption path^16–18^. In the appendicular and axial skeleton, PTH signaling promotes both bone resorption (primarily through increased RANKL expression by mesenchymal lineage cells) and bone formation. PTH and PTHrP play important roles in balancing bone production and bone resorption, with varying effects depending on the quantity and frequency administered^15,19,20^. Osteo-anabolic actions of PTH include suppressing osteoblast apoptosis, reducing osteocyte expression of osteoblast inhibitors, releasing osteogenic factors from resorbed bone matrix, enhancing osteoblast progenitor differentiation, and impacting immune cells in the bone marrow microenvironment^14,21^. Intermittent PTH administration stimulates bone formation and increases bone mass in appendicular and axial skeletal tissues and craniofacial bones^22^.

PTH1R, a G protein-coupled receptor (GPCR), is activated by PTH and PTHrP, leading to increased intracellular production of cAMP and other secondary messengers^23^. Salt inducible kinases (SIKs), a subfamily of AMP-activated protein kinase (AMPK) family kinases, are downstream targets of PTH signaling in long bones and kidneys^24–28^. PTH/PTHrP signaling triggers PKA-dependent phosphorylation and allosteric inactivation of SIKs. Upon SIK inhibition, phosphorylation of their substrates, such as class IIa HDAC and CRTC proteins, is decreased, allowing these proteins to translocate to the nucleus and regulate target genes. Nuclear class IIa HDACs mainly restrict MEF2-driven gene expression, while nuclear CRTC proteins coactivate CREB-regulated target genes^24–26,29^. In osteocytes, the MEF2 family transcription factor controls sclerostin expression, and the CREB-related transcription factors stimulate RANKL gene expression. HDAC4/HDAC5 are important for PTH-mediated inactivation of sclerostin, which enhances bone formation, while CRTCs are key for upregulating RANKL, thereby promoting bone resorption^27,30^. SIK2 and SIK3 combined deletion, globally and specifically in osteoblasts/osteocytes, leads to increased trabecular bone mass and turnover in the axial and appendicular skeleton. However, these genetic SIK modifications also result in increased cortical bone porosity and reduced cortical bone mass^24,27^.

The function of SIKs in teeth and the craniofacial skeleton remains unknown. Here, we report that SIK2 and SIK3 play crucial roles in alveolar bone development, maintenance, and healing. Surprisingly, unlike in long bones where SIK gene deletion enhances osteoblastic bone formation and increases cancellous bone mass, global deletion of SIK2 and SIK3 in developing and adult mice leads to loss of alveolar mineralized tissue due to impaired bone formation. SIK2/SIK3 gene deletion led to an expansion of phenotypically immature mesenchymal cells in the alveolar bone area, suggesting that SIK2 and SIK3 are essential for appropriate osteoblast maturation during alveolar bone growth and remodeling. The lack of SIKs adversely impacts alveolar bone homeostasis in skeletally mature mice and tooth socket healing after tooth extraction, resulting in the cessation of the final osteoblast maturation. The pronounced phenotypic differences between long bones and alveolar bones after SIK2/SIK3 deletion led us to investigate mechanisms underlying these distinct outcomes. Analysis of single-cell RNA sequencing (scRNA-seq) data from osteoblasts originating from both bone types uncovered transcriptomic differences, which may account for dramatic differences in consequences of SIK2/SIK3 deletion from different groups of osteoblasts.

## Results

### Absent alveolar bone following genetic deletion of SIK2/SIK in developing mice

We utilized genetic models to study the role of Salt Inducible Kinases in alveolar bone development. Of the three mammalian SIK isoforms, SIK2 and SIK3 (genes closely linked to syntenic regions of mouse chromosome 9 and human chromosome 11) are important downstream regulators of PTH1R action in bone and kidney. Therefore, first we used *Sik2*^f/f^*; Sik3*^f/f^*; ubiquitin-*Cre^ERt2^ (mutant) mice^31^ to achieve inducible/global SIK2 and SIK3 deletion. To induce deletion, tamoxifen was given to the mutant and control (*Sik2*^f/f^*; Sik3*^f/f^) at postnatal day 3 (P3) when the tooth roots of mandibular molars start to form. Mandibles were then examined at P8, P11, P14, P21 and P28 **(Figure 1A-J)**. At P8, no significant differences were seen between control and mutant mandibles. Surprisingly, we noted decreased alveolar bone in the SIK2/SIK3 double knockout mandible as early as P11. By P21, SIK2/SIK3 mutant mandibles showed essentially no alveolar bone in both inter-radicular and inter-septal bone regions, with fibrous-like tissue replacing the area. Long bones (femurs) at P28 from mice treated with tamoxifen at P3 exhibited the expected changes from SIK2/SIK3 inhibition, including increased trabecular bone and expanded proliferating chondrocytes with substantial truncation of bone length **(Figure 1K, L)**^27,32^.

**Figure 1:**
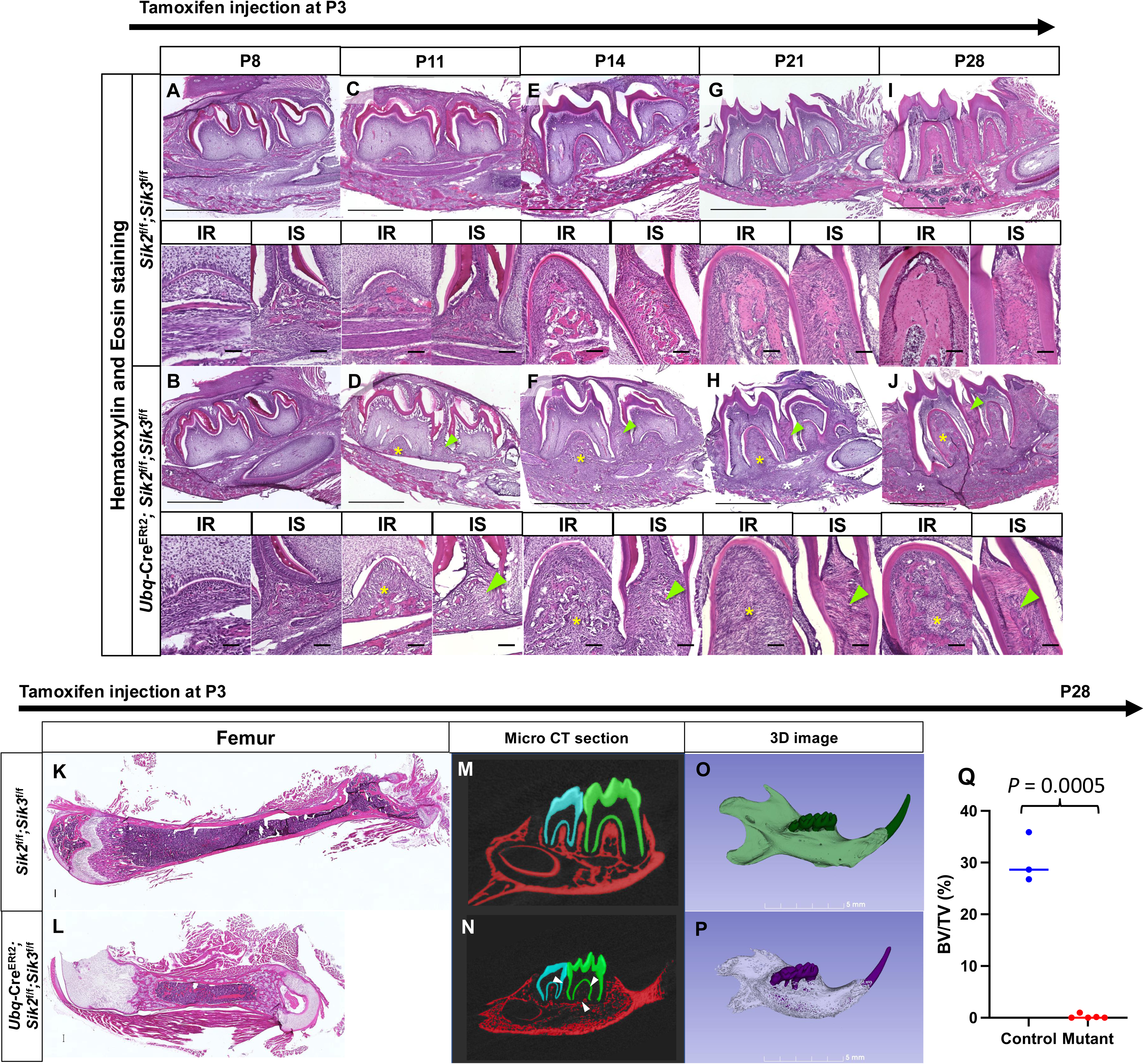
Reduced alveolar bone following global SIK2/SIK3 deletion at postnatal day 3. **(A, C, E, G, I)** H&E staining of the control mandible (*Sik2*^f/f^ *; Sik3*^f/f^) at P8, P11, P14, P21 and P28 respectively. Upper panel: 10x mandible (Scale bar: 1000 μm), Lower panel: 20x interradicular and interseptal bone (Scale bar: 100 μm) **(B, D, F, H, J)** H&E staining of the mutant mandible (*ubiquitin-*Cre^ERt2^ *; Sik2*f/f*;Sik3*^f/f^) at P8, P11, P14, P21, and P28 respectively. Upper panel: 10x mandible (Scale bar: 1000 μm), Lower panel: 20x interradicular and interseptal bone (Scale bar: 100 μm) Yellow asterisk: Reduced interradicular bone in the mutant mandibles. Green arrowhead: Reduced interseptal bone in the mutant mandibles. IS: Interseptal bone IR: Interradicular bone **(K, L)** H&E staining of P28 control and mutant femur, respectively. Scale bar: 1000 μm **(M, N)** Sagittal section of Micro CT of the control and the mutant mandible, White arrowhead: Punched-out lesion in alveolar bone area. **(O, P)** Three dimensional rendering images of the control and the mutant mandible **(Q)** BV/TV of the first mandibular interradicular bone area of the control (blue) and the mutant (red) Students t-test was performed p-value < 0.05

To further define changes in alveolar bone mineralization, micro-CT was performed on P28 mandibles following P3 tamoxifen administration **(Figure 1M-P)**. Mutant mandibles were smaller in size and expanded in the sagittal plane with increased cortical bone porosity. The most noticeable characteristic of the mutant mandibles was a “punched-out” lesion surrounding the molars, consistent with the absence of alveolar bone that was noted histologically. In mutant mandibles, the alveolar bone volume fraction was considerably reduced **(Figure 1Q).** Overall, histology and micro-CT analysis of mandibles at P28 revealed an unexpected loss of alveolar bone without obvious alterations in tooth morphology or eruption following SIK2/SIK3 deletion at P3.

### Periodontial expression of SIK isoforms

Based on observed alveolar bone defects in mice with SIK2/SIK3 ablation, we next surveyed SIK isoform expression by RNAscope in P10 mandible sections. *Sik1* was primarily expressed in the periodontal ligament cells in interradicular and interseptal bone areas **(Figure 2A-D)**. *Sik2* was found in periodontal ligament cells and alveolar bone osteocytes **(Figure 2E-H)**. *Sik3* expression was qualitatively highest in PDL cells and alveolar bone osteoblasts and osteocytes; furthermore, *Sik2* and *Sik3* expression was noted in the cortical bone osteocytes and periosteum at the inferior border of the mandible **(Figure 2I-L)**. Thus, SIK isoforms, particularly *Sik2* and *Sik3*, are expressed locally in cells participating in alveolar bone formation and remodeling.

**Figure 2:**
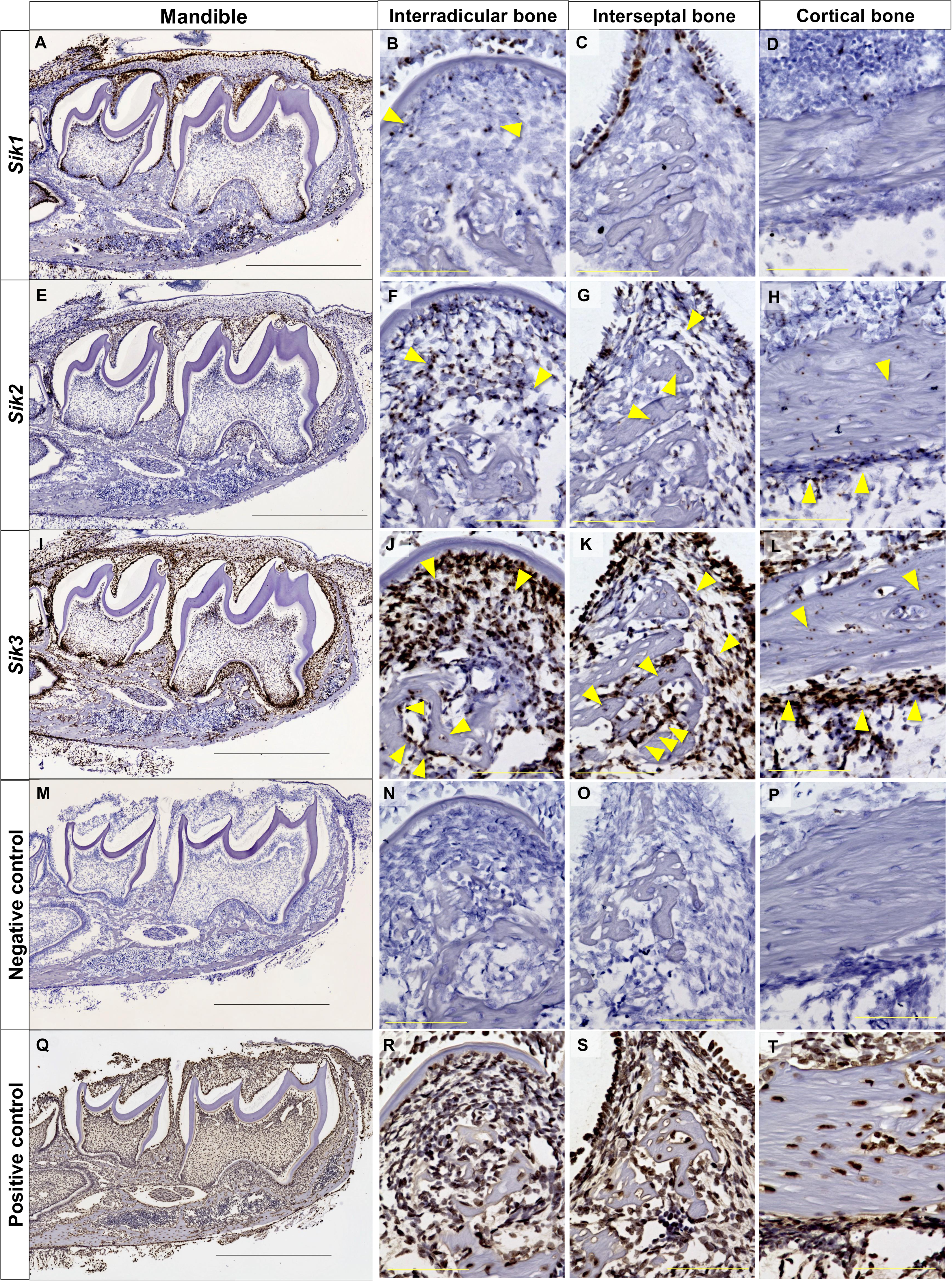
Periodontal SIK isoform expression. **(A, B, C, D)** *Sik1* in situ hybridization (brown, RNAscope) in paraffin-section of mandible (10x), interradicular bone (40x), interseptal bone (40x), and mandibular cortical bone (40x) at P10. Yellow arrowheads: *Sik1* expression in periodontal ligaments. **(E, F, G, H)** *Sik2* in situ hybridization (brown, RNAscope) in paraffin-section of mandible (10x), interradicular bone (40x), interseptal bone (40x), and mandibular cortical bone (40x) at P10. Yellow arrowheads: *Sik2* expression in periodontal ligaments and osteocytes. **(I, J, K, L)** *Sik3* in situ hybridization (brown, RNAscope) in paraffin-section of mandible (10x), interradicular bone (40x), interseptal bone (40x), and mandibular cortical bone (40x) at P10. Yellow arrowheads: *Sik3* expression in periodontal ligament, osteoblast, osteocytes and periosteal of mandible. **(M, N, O, P)** Positive control in situ hybridization (brown, RNAscope) in paraffin-section of mandible (10x), interradicular bone (40x), interseptal bone (40x), and mandibular cortical bone (40x) at P10. **(Q, R, S, T)** Negative control in situ hybridization (brown, RNAscope) in paraffin-section of mandible (10x), interradicular bone (40x), interseptal bone (40x), and mandibular cortical bone (40x) at P10. Scale bar: 1000 μm for 10x and 100 μm for 40x

### Global SIK2/SIK3 deletion disrupts alveolar formation during jaw development

We then asked whether alveolar bone defects were due to decreased bone formation or increased bone resorption. In long bone, SIK2/SIK3 deletion increases RANKL expression and osteoclast activity^31^. Therefore, first we performed TRAP staining to assess osteoclast numbers **(Figure 3A-N)**. Surprisingly, at P8 and P11, there was no discernible difference in TRAP staining patterns between the two genotypes. Notably, and contrary to our expectations, starting at P14, a significant *reduction* of osteoclasts per tissue area was observed in the interseptal bone area of SIK2/SIK3 mutant mice. By P28, little to no TRAP-positive cells were observed in the interseptal and interradicular bone area **(Figure 3O)**. Thus, enhanced osteoclastic bone resorption is not the primary cause of alveolar bone loss in this model. Von Kossa staining on non-decalcified sections was performed to visualize mineralized tissue of the alveolar bone area at P14. Less mineralization tissue was observed in the mutant mandible compared to the control **(Figure 3P-R: Left panels)**. Next, we performed *in vivo* fluorochrome labeling to visualize bone deposition. Calcein was injected twice at P10 and P12, and mandibles were dissected at P14 for non-decalcified histology. Notably, SIK2/SIK3 mutants showed reduced calcein deposition in the alveolar bone area compared to controls **(Figure 3P-R: middle and right panels)**. Taken together, these findings show that loss of alveolar bone in global/inducible SIK2/SIK3 mutants is likely due to defective alveolar bone formation rather than increased bone resorption.

**Figure 3:**
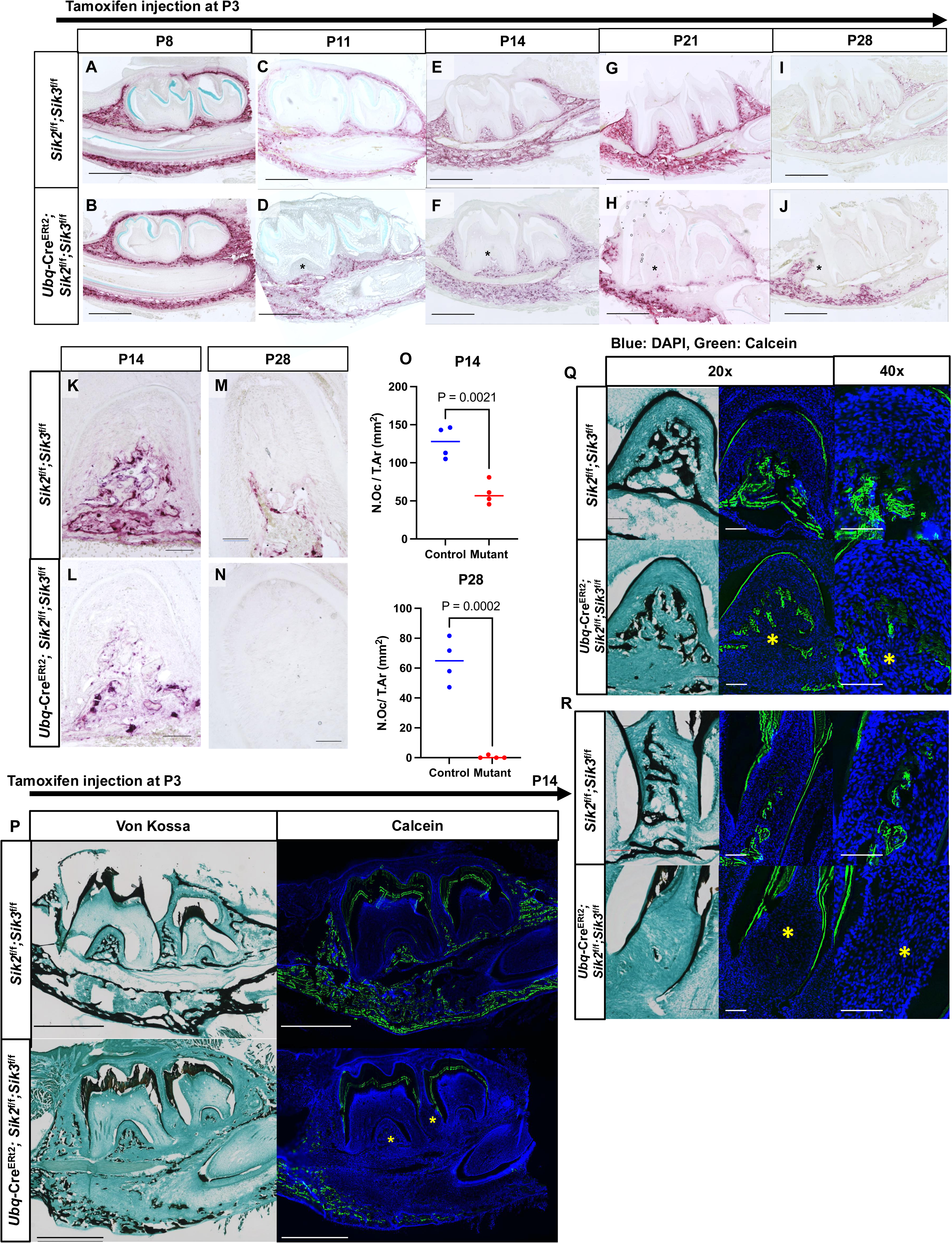
Reduced alveolar bone formation in global SIK2/SIK3 mutant mice. **(A, C, E, G, I)** TRAP staining of the control mandible at P8, P11, P14, P21 and P28 respectively. Scale bar: 1000 μm **(B, D, F, H, J)** TRAP staining of the mutant mandible (*ubiquitin-*Cre^ERt2^ *; Sik2*^f/f^ *; Sik3*^f/f^) at P8, P11, P14, P21, and P28 respectively. Black asterisk: Decreased osteoclast number in alveolar bone. Scale bar: 1000 μm **(K, M)** 20x TRAP staining of control mandible at P14 and P28, respectively. Scale bar: 100 μm **(L, N)** 20x TRAP staining of mutant mandible at P14 and P28, respectively. Scale bar: 100 μm **(O)** Number of Osteoclast per area of control (Blue) and mutant (Red) at P14 (upper) and P28 (lower) Students t-test was performed p-value < 0.05 **(P)** Von Kossa staining (Left) and calcein labeling (Right) of the uncalcified mandible at P14. Scale bar: 100 μm **(Q)** Von Kossa staining (Left) and calcein labeling (Right) of interradicular area at P14. Scale bar: 100 μm **(R)** Von Kossa staining (Left) and calcein labeling (Right) of interseptal area at P14. Scale bar: 100 μm Yellow asterisk: Decreased calcein labeling in alveolar bone area

SIK2/SIK3 deletion with *ubiquitin-*Cre*^ERt2^* in long bones causes substantial increases in trabecular bone mass due to increased bone formation^31^. Therefore, noting *reduced* alveolar bone formation in this same model was surprising. To investigate this unexpected observation, we assessed osteoblast maturation by staining control and mutant alveolar bones at P28 with Alkaline phosphatase (Alk Phos) and Osteocalcin (Ocn), markers of early and mature osteoblasts, respectively **(Figure 4A-H)**. Despite reduced mineralized tissue and calcein deposition, we noted statistically significant increases in alkaline phosphatase expression in mutant mandibles. However, these alkaline phosphatase-positive cells interspersed throughout regions of absent alveolar bone did not express osteocalcin, indicating that SIK2/SIK3 loss impairs alveolar bone osteoblast terminal maturation.

**Figure 4:**
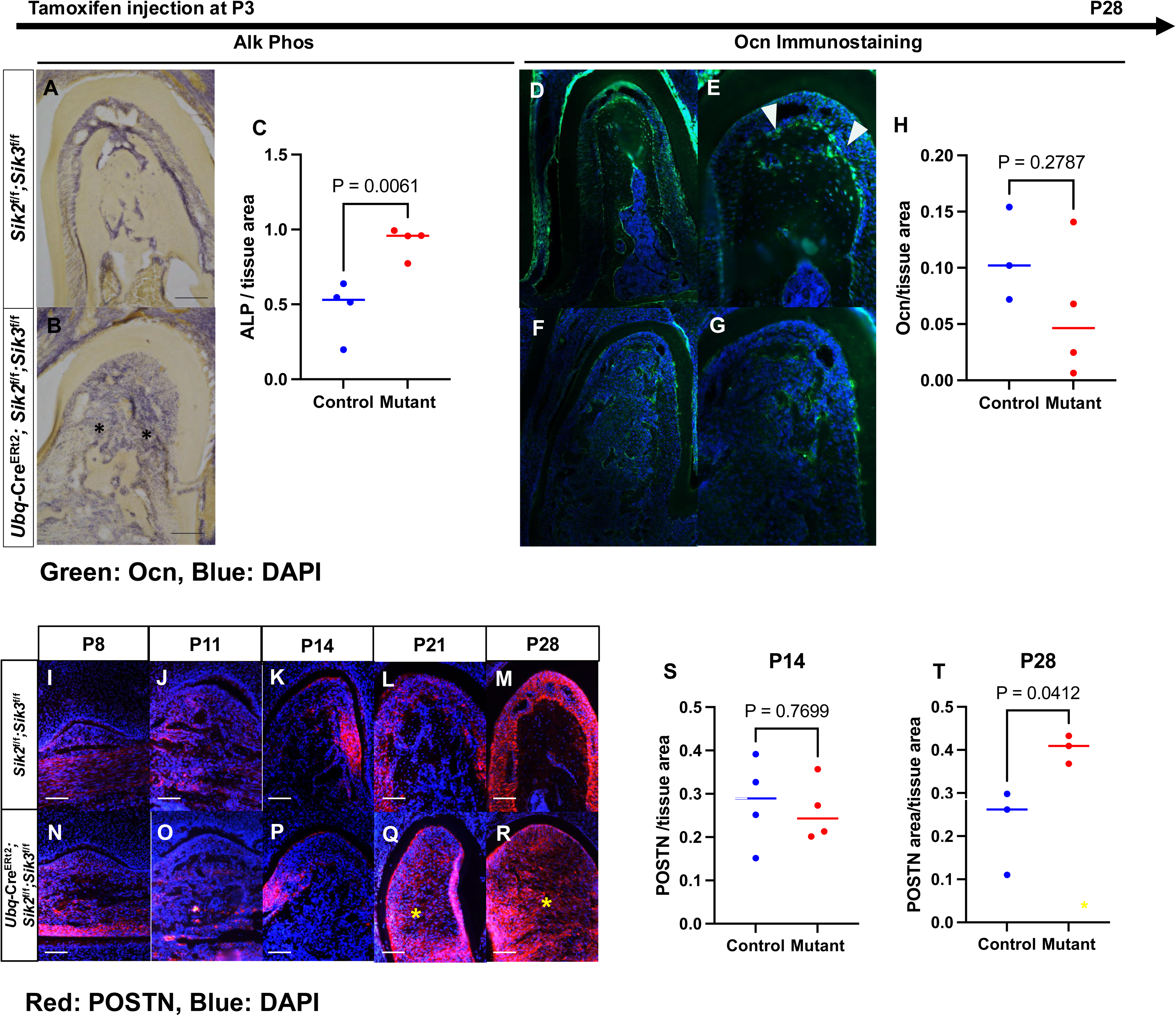
Arrested osteoblast maturation with increased Periostin expression in global SIK2/SIK3 mutant mice. **(A)** Alkaline phosphatase staining of the control mandible in the interradicular bone area. **(B)** Alkaline phosphatase staining of the mutant mandible in the interradicular bone area. Black asterisk: bone loss area with Alkaline phosphatase positive cells. **(C)** Alkaline phosphatase positive per tissue area of control (Blue) and mutant (Red) at P28. Students t-test was performed p-value < 0.05 **(D, E)** Ocn immunostaining of control mandible in interseptal bone area, 10x and 20x, respectively. White arrowhead: Osteocalcin positive osteoblasts **(F, G)** Ocn immunostaining of the mutant mandible in interradicular bone area, 10x and 20x, respectively. **(H)** Ocn expression per tissue area of control (Blue) and mutant (Red) at P28. Students t-test was performed p-value < 0.05 **(I-M)** Periostin immunostaining of control mandible at interradicular area from P8, P11, P14, P21, and P28 respectively. **(N-R)** Periostin immunostaining of *ubiquitin-*Cre^ERt2^ *; Sik2*^f/f^ *; Sik3*^f/f^ at interradicular area from P8, P11, P14, P21 and P28 respectively Scale bar: 100 μm Yellow asterisk: bone loss area with Periostin-positive cells **(S-T)** Periostin positive area per tissue area of control (Blue) and mutant (Red) at P28. Students t-test was performed p-value < 0.05

PTH1R deletion in PTHrP-positive dental follicle cells causes a shift in mesenchymal cell fate from PDL cells and acellular cementoblasts to cementoblast-like cells, impairing PDL differentiation and leading to failed tooth eruption^9^. Since PTH1R signaling suppresses cellular SIK2/SIK3 activity, we considered the possibility that SIK2/SIK3 deletion could result in a phenotype opposite that of PTH1R deficiency. Importantly, increased Periostin (POSTN, a marker of PDL fibroblasts^33^) levels are seen in the PDL of PTH1R activation models^10^. Therefore, we performed anti-POSTN immunostaining at all time points **(Figure 4M-V)**. There were no discernible changes between the mutant and control mandibles at P14. However, at P21 and P28, significantly increased POSTN-positive cells were observed in inter-septal and inter-radicular areas in areas with absent alveolar bone in SIK2/SIK3 mutants. This finding suggests that the absence of SIK2 and SIK3 impacts mesenchymal cell development and causes a population shift from mature bone-forming osteoblasts to ectopic POSTN-positive cells.

### SIK2/SIK3 are important for maintaining alveolar bone in adult mice

Having established a key role for SIK2/SIK3 in alveolar bone development using P3 tamoxifen injections, we next asked if SIKs contribute to alveolar bone homeostasis in skeletally mature mice. Therefore, we globally ablated SIK2/SIK3 in 12-week-old mice using *ubiquitin*-Cre^ERt2^. 2 weeks after tamoxifen injection, no significant alveolar bone differences were observed between control and knockout littermates. However, by 8 and 10 weeks following SIK2/SIK3 deletion, loss of alveolar bone was noted in mutant mice in both interradicular and interseptal areas **(Figure 5A-F)**.

**Figure 5:**
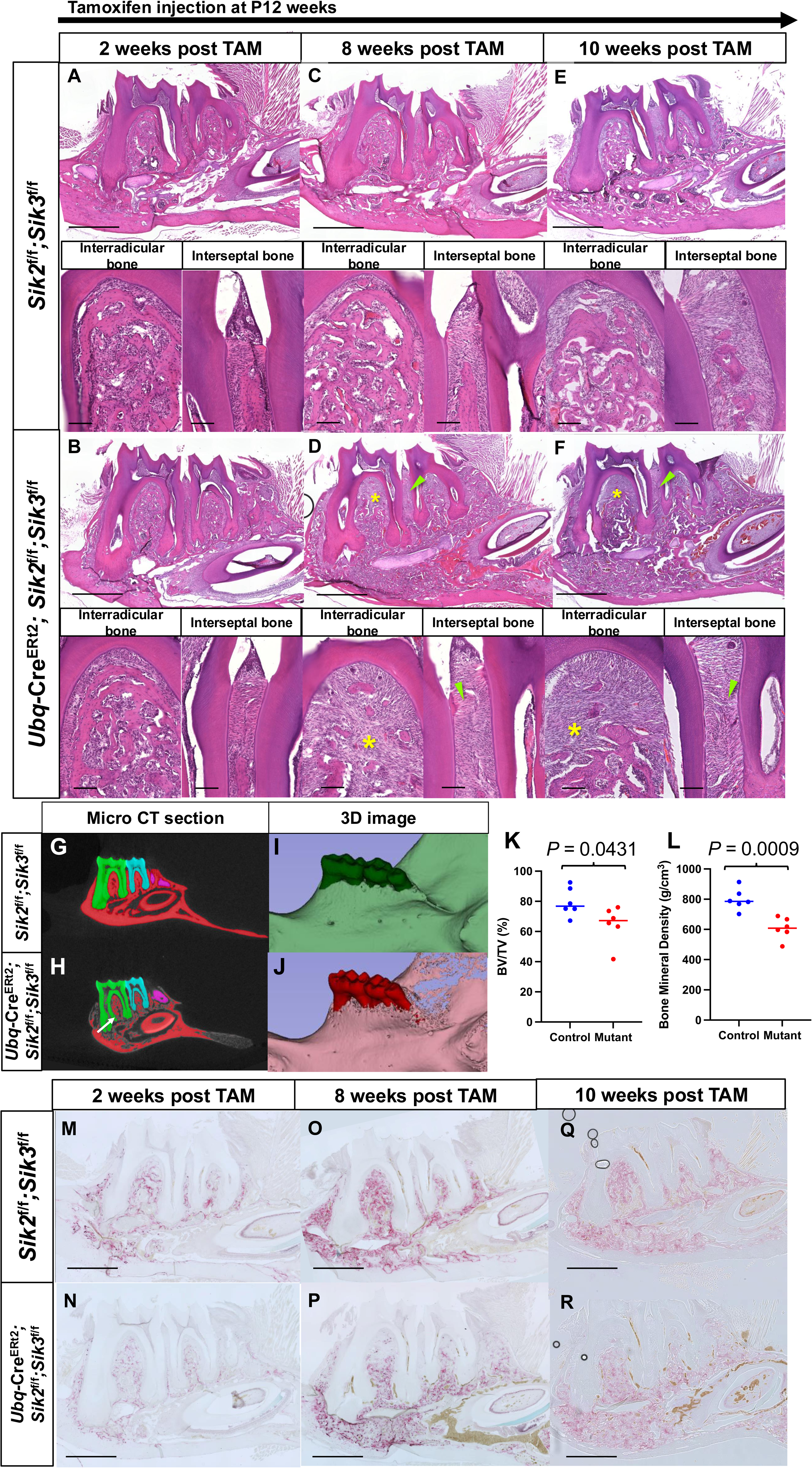
Global deletion of SIK2/SIK3 in adult mice induces alveolar bone loss. **(A, C, E)** H&E staining of the control mandible 2 weeks, 8 weeks, and 10 weeks (respectively) after tamoxifen injection. Upper panel: 10x mandible (Scale bar: 1000 μm), Lower panel: 20x interradicular and interseptal bone (Scale bar: 100 μm) **(B, D, F)** H&E staining of mutant mandible 2 weeks, 8 weeks, and 10 weeks (respectively) after tamoxifen injection. Yellow asterisk: Interradicular bone loss, Green arrowhead: Interseptal bone loss Upper panel: 10x mandible (Scale bar: 1000 μm), Lower panel: 20x interradicular and interseptal bone (Scale bar: 100 μm) **(G, I)** Micro CT section and 3D rendering of 10 weeks post-tamoxifen injected control mandible. **(H, J)** Micro CT section and 3D rendering of 10 weeks post-tamoxifen injected mutant mandible. **(K)** Bone volume fraction of control and mutant first mandibular molar interradicular bone area. Students t-test was performed p-value < 0.05 **(L)** Bone mineral density of control and mutant first mandibular molar interradicular bone area. Blue: Control, Red: Mutant Students t-test was performed p-value < 0.05 **(M, O, Q)** TRAP staining of the control mandible 2 weeks, 8 weeks, and 10 weeks (respectively) after tamoxifen injection. Scale bar: 1000 μm **(N, P, R)** TRAP staining of *ubiquitin-*Cre^ERt2^ *; Sik2*^flf^ *; Sik3*^f/f^ mandible 2 weeks, 8 weeks, and 10 weeks (respectively) after tamoxifen injection. Black asterisk: Less osteoclast number in the alveolar bone. Scale bar: 1000 μm

To further explore changes from SIK2/SIK3 deletion in mature alveolar bone tissue, we conducted micro-CT analysis in mandibles 10 weeks post-tamoxifen injection **(Figure 4G-J)**. The mutant mandibles showed reduced alveolar bone mineral density and alveolar bone volume fraction **(Figure 5K-L)**, findings in stark contrast to the long bone phenotype (increased trabecular bone mass) reported in these same mice^31^. Mutant alveolar bone regions showed decreased TRAP staining **(Figure 5M-R)**, findings consistent with phenotypes observed in the developmental model. In sum, these data again indicate that decreased bone formation, not increased bone resorption, is most likely to cause of alveolar bone loss in this model. These data show that SIK2 and SIK3 are important for both alveolar bone development and maintenance.

### SIK2/SIK3 inhibition impaired tooth socket healing

Tooth extraction initiates a cascade of healing processes within the alveolar socket, involving a delicate interaction between bone resorption and new bone formation. Tooth socket healing illustrates the intricate relationship between developmental processes and regenerative capacity. Given our previous result demonstrating that SIK2/SIK3 play crucial roles in alveolar bone osteoblast differentiation, we explored how inhibiting SIK2/SIK3 affects alveolar bone regeneration. Control and mutant mice were treated with tamoxifen at 12 weeks of age, 10 weeks later, the timepoint when alveolar bone defects are noted (**Figure 5E-F**), extraction of the first maxillary molar was performed, followed by analysis 4 weeks later. At this time, control maxilla exhibited completed healing with organized epithelium and mineralized bone in the extraction sockets. In contrast, mutant maxilla showed severe alveolar bone loss with extraction sockets filled with fibrous-like tissue **(Figure 6A-D).** Micro-CT analysis of the SIK2/SIK3 deleted maxilla confirmed the absence of mineralization within the tooth socket, with the bone volume fraction in this region reduced to almost zero **(Figure 6E-G)**.

**Figure 6:**
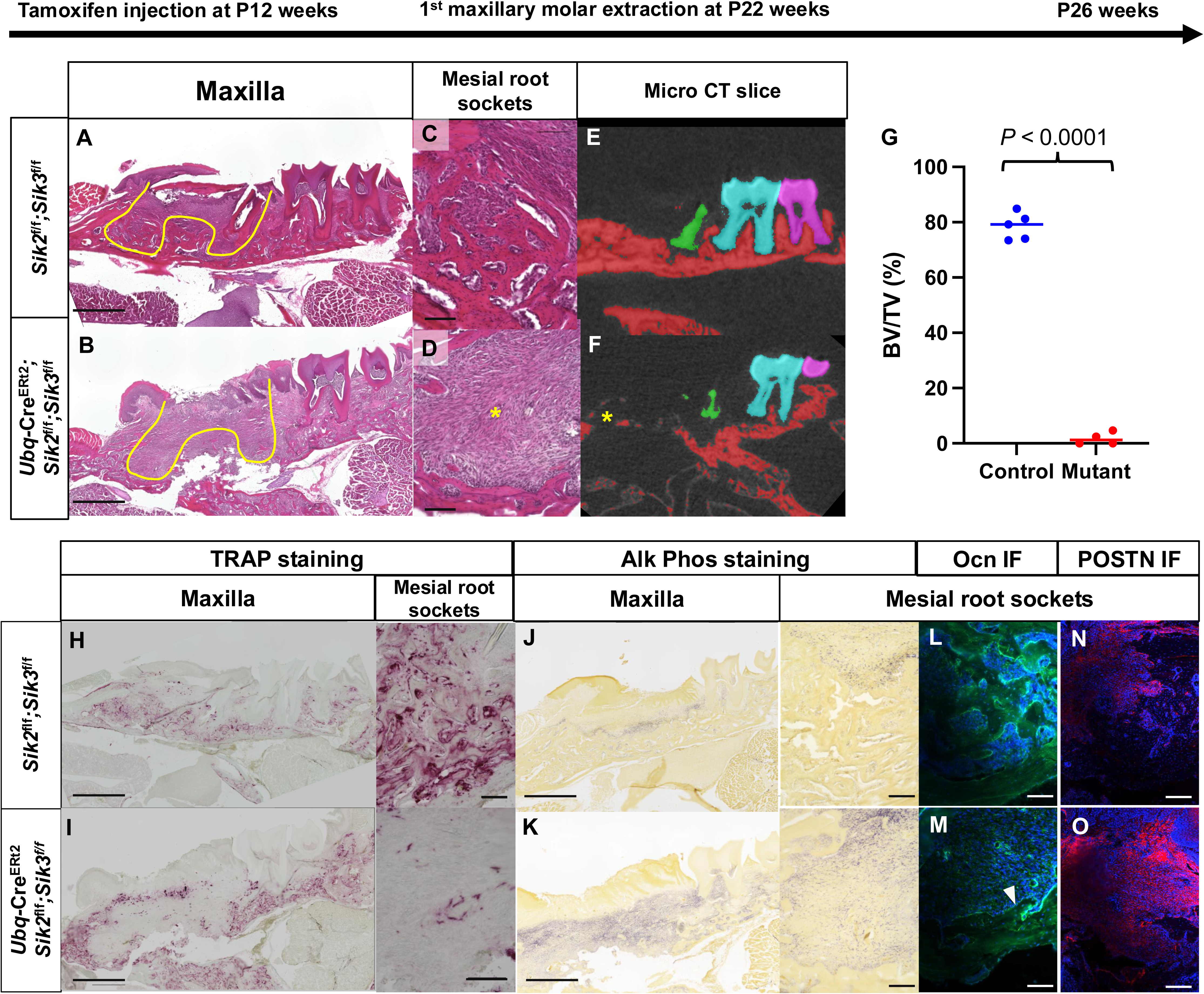
Impaired socket healing after tooth extraction in SIK2/SIK3 mutant mice. **(A, B)** H&E staining of the control and mutant maxilla, respectively. Scale bar: 1000 μm **(C, D)** 20x magnification of H&E staining of control and mutant at mesial root socket, respectively. Scale bar: 100 μm **(E, F)** Micro CT sagittal sections of micro-CT of the control and mutant maxilla respectively. Yellow asterisk: Lack of bone formation in tooth socket **(G)** Bone volume fraction of first maxillary molar sockets of control (Blue) and mutant (Red). Students t-test was performed p-value < 0.05 **(H, I)** TRAP staining of control and mutant maxilla and mesial root sockets. Scale bar: 1000 μm (Right panel), 100 μm (Left panel) **(J, K)** Alkaline phosphatase staining of control and mutant maxilla and mesial root sockets. Scale bar: 1000 μm (Right panel), 100 μm (Left panel) **(L, M)** Osteocalcin Immunofluorescent in mesial root sockets of the control and mutant maxilla, respectively. White arrowhead: less osteocalcin staining at the bottom of the mesial root socket. Scale bar: 100 μm **(N, O)** Periostin Immunofluorescent in mesial root sockets of control and mutant maxilla, respectively. Scale bar: 100 μm

Reduced TRAP staining in mesial root sockets indicates that the tooth socket healing defects are not due to increased bone resorption **(Figure 6H-I)**. To assess osteoblast activity, alkaline phosphatase staining, and osteocalcin immunostaining were performed. Alkaline phosphatase expression was increased and distributed throughout the mutant tooth socket, while osteocalcin expression was reduced in the extraction socket floor of the mutant mice **(Figure 6J-M)**. These findings are consistent with data in our developmental model. Additionally, we conducted anti-periostin immunostaining to characterize the fibrous-like tissue that replaces bone during tooth socket healing **(Figure 6N-O)**. We noted an upregulation of periostin-positive fibrous tissue, suggesting an expansion of ectopic periodontal ligament (PDL)-like cells in mutant tooth sockets. In conclusion, these results suggest that SIK2 and SIK3 are important for proper tooth extraction healing where they play an important role in the maturation of alveolar bone osteoblasts.

### Osteoblasts from long bone and alveolar bone show different transcriptomic profiles

The striking phenotypic difference observed between long bones and alveolar bones following SIK2/SIK3 deletion prompted us to explore mechanisms responsible for these distinct outcomes. To this end, we compared previously reported single-cell RNA sequencing (scRNA-seq) datasets from long and alveolar bone osteoblasts. scRNA-seq profiles from long bone mesenchymal cells marked by Prrx1-cre; R26RtdTomato at P21^34^ and alveolar bone osteoblasts marked by Col1(2.3kb)-GFP at P25^35^ were computationally merged with Harmony^36^ **(Figure 7A-B)**. Unsupervised clustering of the merged dataset revealed a total of 18 cell clusters **(Figure 7C)**. Among these clusters, clusters 0, 1, 2, 14, and 16 exhibited the highest expression of osteoblast marker genes (*Col1a1*, *Bglap*, *Sp7*, and *Dmp1*) indicating that these osteoblast populations were common in both datasets **(Figure 7D)**. To identify the transcriptomic differences between osteoblasts from long bone and alveolar bone, cluster-specific differential expression gene (DEG) analysis was performed on these osteoblastic cell clusters. As expected, based on the genetic tools used to isolate these cells, tdTomato transgene expression was significantly increased in osteoblasts from long bone, whereas alveolar bone osteoblasts showed increased GFP transgene expression. Furthermore, the majority of the osteoblast clusters (clusters 0, 1, 2, and 16) showed a considerable number of DEGs based on the site of isolation **(Figure 7E)**. Most notably, cluster 1 had a total of 1467 DEGs (402 DEGs upregulated in long bone and 1065 DEGs upregulated in alveolar bone). Gene ontology (GO) enrichment analysis of DEGs in cluster 1 revealed significant enrichment of several GO terms in alveolar bone, including extracellular structure organization (GO:0043062), axon guidance (GO:0007411) and embryonic skeletal system morphogenesis (GO:0048704) **(Figure 7G)**. Despite these transcriptomic differences, expression of *Sik2* and *Sik3* was not significantly different between osteoblasts of different anatomic origin **(Figure 7F)**.

**Figure 7:**
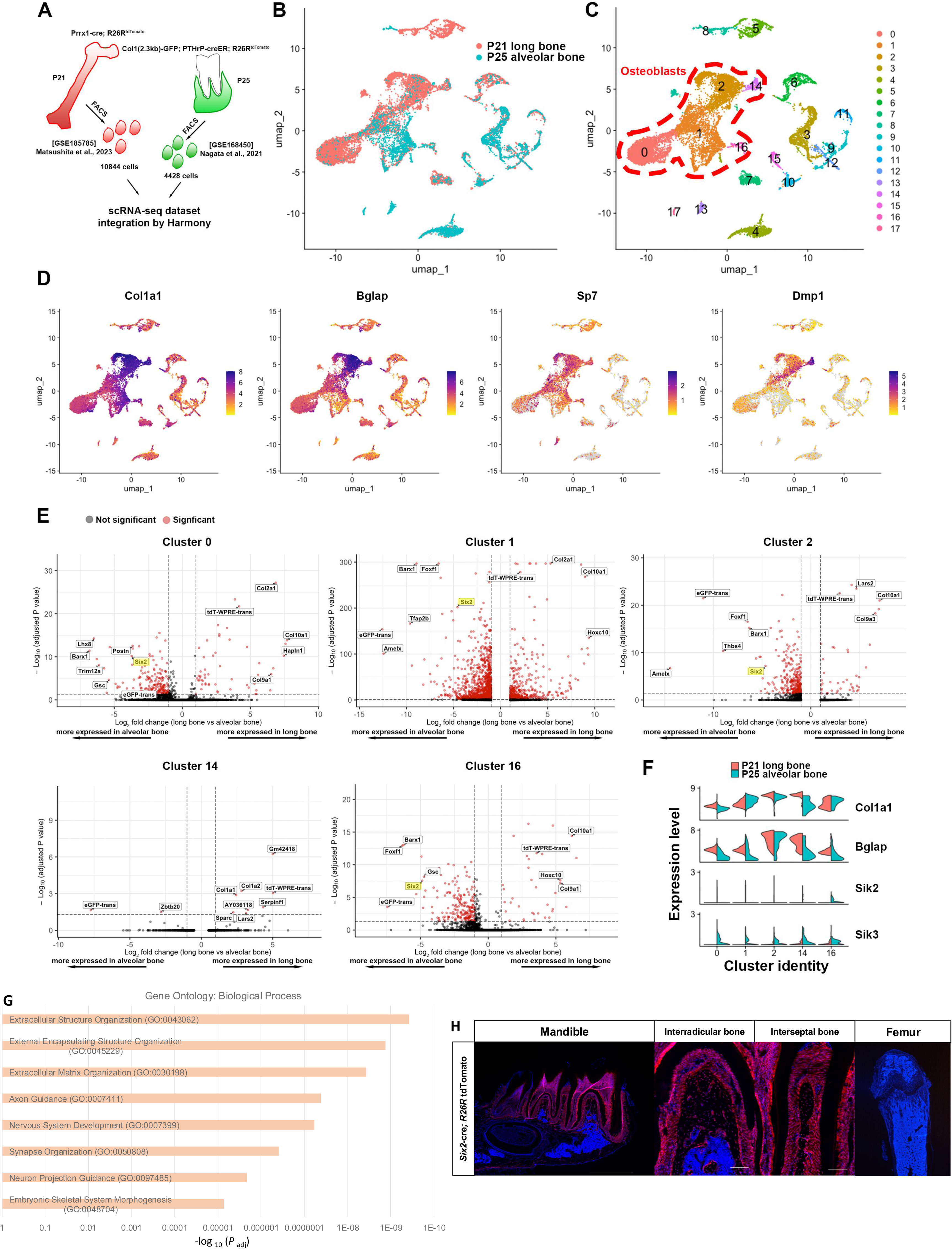
Single cell transcriptome analysis of long bone and alveolar bone osteoblasts. **(A)** Schematic for scRNA-seq workflow. Datasets of sorted tdTomato+ cells isolated from *Prrx1*-cre; *R26R*tdTomato femurs at P21 [GSE185785] and Col1-GFP+ cells from *Col1(2.3kb)-GFP; PTHrP-*cre^ER^; *R26R*tdTomato alveolar bones at P25 [GSE168450] were integrated using Harmony. **(B)** UMAP visualization of the two datasets (P21 femur long bone and P25 alveolar bone) merged by Harmony. Each dot represents an individual cell. **(C)** UMAP visualization of major cell clusters. Red dotted outline: osteoblast clusters. **(D)** FeaturePlots of osteoblast specific markers. High gene expression: violet, low expression: yellow, no expression: grey. **(E)** Volcano plots of DEGs from P21 long bone versus P25 alveolar bone osteoblast clusters. DEGs indicated by red dot plots are considered significant by adjusted P value < 0.05 and |average log2 fold change| > 1. **(F)** Violin plots show normalized gene expression level in each down sampled osteoblast cell clusters. **(G)** Gene ontology enrichment analysis of DEGs between P25 alveolar bone and P21 long bone. **(H)** Tomato expression of *Six2*-cre; *R26R-*tdTomato in mandible of P25 mouse. Scale bar: 1000 μm (Mandible, Femur), 100 μm (Interradicular bone, Interseptal bone)

Further analysis of genes enriched in alveolar bone osteoblasts revealed that the transcription factor SIX2, a gene that is highly expressed in neural crest-derived mesenchymal cells^37,38^, is expressed in alveolar bone but not in long bone. Therefore, we evaluated *SIX2* expression in alveolar bone in *Six2*-cre; *R26R*-tdTomato mice^39^. At P28, we observed abundant tdTomato+ cells in alveolar bone osteoblasts, osteocytes, periodontal ligament, and in the pulpal area, while no tdTomato+ cells were observed in long bone^28^ **(Figure 7H)**. Taken together, these data indicate that alveolar bone osteoblasts show transcriptomic differences when compared to long bone osteoblasts and that SIX2-Cre may be a useful strain to selectively ablate genes in this osteoblast subset.

## Discussion

Our results suggest that SIK2/SIK3 are important for normal alveolar bone development and alveolar bone homeostasis/remodeling. Alveolar bone defects in SIK2/SIK3 mutant mice are associated with poor tooth socket healing following molar extraction. In all models, TRAP staining was reduced in SIK2/SIK3 mutant mice, indicating that decreased bone formation rather than increased bone resorption is the primary cause of alveolar bone loss. Surprisingly, in both developmental and tooth extraction models, we noted increased alkaline phosphatase expression upon SIK2/SIK3 deletion without upregulation of the mature osteoblast marker osteocalcin. Periostin-positive fibrous tissue was present in zones of absent alveolar bone in developmental and post-extraction models. This phenotype resembles what is seen in continuous hyperparathyroidism, a condition associated with alveolar bone loss. Patients with hyperparathyroidism exhibit reduced alveolar bone density and loss of lamina dura with extension of periodontal ligament space around the tooth root^40–43^. Additionally, there is a correlation between the degree of periodontal ligament expansion and the degree of PTH elevation in blood^41,42^. Furthermore, mandibular cortical bone density is reduced in hyperparathyroidism^41,44^. However, it is thought that alveolar bone defects linked to hyperparathyroidism are mainly due to increased bone turnover and bone resorption^45,46^. In contrast, our findings suggest that the primary issue in the absence of SIK2/SIK3 is reduced matrix production and mineralization by mature alveolar bone osteoblasts. Further study is necessary to elucidate the molecular mechanisms by which SIK2/SIK3 selectively regulates osteoblast maturation in alveolar bone.

One limitation of our study is that we globally eliminated SIKs in all tissues using the broadly expressed *ubiquitin-*Cre^ERt2^ driver. Thus, at present, we cannot identify the specific cell type(s) in which SIK2/SIK3 deletion causes alveolar bone defects. For instance, it remains possible that systemic mineral metabolism parameters, such as mild hypercalcemia or modestly elevated 1,25-vitamin D levels observed in global SIK2/SIK3 mutants^28^, might indirectly influence alveolar bone development and remodeling^47^. Previous studies have shown that PTHrP signaling is crucial for both tooth and alveolar bone development^9,11,48,49^. Despite their anatomical proximity, our mouse model did not exhibit any significant tooth phenotype, suggesting that the deletion of SIK2/3 does not critically affect tooth development. This finding implies that the alveolar bone loss observed in our study is unlikely to be a consequence of changes in the tooth due to SIK2/3 deletion. However, further research is necessary to determine whether the alveolar bone phenotype arises directly from bone cells or from interactions between alveolar bone cells and the periodontal ligament. Thus, although our study offers insight into the role of SIKs in alveolar bone, it cannot conclusively establish a cell-intrinsic role of these kinases in alveolar bone osteoblast progenitors. Future studies using cell type-specific Cre drivers, such as *Six2*-Cre^28^, to delete SIK2/SIK3 will likely provide further insights into this important question. Given the cell-intrinsic effects of SIK2/SIK3 gene deletion in long bone seen using *Dmp1*-Cre^27^ and the broad expression of these kinases in craniofacial tissue, it is most likely that a cell-intrinsic role for SIKs in alveolar bone osteoblasts and their progenitors explains the phenotypes observed here.

Another limitation of our study is that we extracted the first maxillary molar 10-weeks after SIK2/SIK3 deletion in adult mice, a time point in which alveolar bone defects were already present. The socket healing process involves alveolar bone loss due to bone remodeling, which entails both bone formation and resorption. Alveolar ‘ridge reduction’ following tooth extraction can be expected in all cases; however, the alveolar bone quantity and quality prior to tooth extraction, among other local factors, determine the extent of bone resorption during this remodeling process^50–54^. Therefore, while we conclude that SIK2/SIK3 are important for alveolar bone healing after tooth extraction, it is crucial to note that mildly reduced alveolar bone prior to extraction may worsen bone healing defects. Nonetheless, adult-onset SIK2/SIK3 mutant mice with reduced alveolar bone offer an excellent model to evaluate the relationship between intact alveolar bone tissue and bone healing following tooth extractions.

Our study revealed a striking divergence in the observed phenotypes between long bones and alveolar bone in response to SIK2/SIK3 deletion. While global deletion of SIK2/SIK3 in long bones increases trabecular bone mass, bone formation, and bone turnover in long bones^31^, defects with decreased bone formation are present in alveolar bone of the same mice. This highlights the emerging concept that different bones within the body exhibit unique behavior and responses to SIK2/SIK3 deletion. This divergence could be attributed to unique cell origins and gene regulatory networks in anatomically-distinct osteoblast subsets, a model supported by our integrated scRNA-seq comparison between long bone and alveolar osteoblasts. Consistent with previous studies that conducted comparative transcriptomic analyses, certain genes show increased expression in alveolar bone. Moreover, GO analysis highlighted enrichment of genes associated with extracellular structure organization and skeletal system morphogenesis in alveolar bone^55,56^. Among the top eight Biological Process GO terms enriched in differentially expressed genes (DEGs) in alveolar bone, several involve nervous system development and neuron organization, such as Axon Guidance, Nervous System Development, and Synapse Organization. This may be attributed to the fact that alveolar bone is derived from ecto-mesenchymal neural crest cells^1^. In addition, we found that chondrocytes (Cluster 5) are mainly found in long bones. This is likely because long bone derives from mesenchymal cells through endochondral ossification^57^, whereas alveolar bone forms from ecto-mesenchymal neural crest cells through intramembranous ossification^1^. Additionally, the unique oral microenvironment, mechanical forces, and local signaling factors may further underlie different behaviors between alveolar and long bone osteoblasts. Due to the constant exposure to occlusal stress and its adjacency to oral biofilms, the metabolism and remodeling processes of alveolar bone are the most dynamic in the skeletal system, being three to sixfold more robust than other skeletal tissue^8,58–61^. Therefore, our study emphasizes the need to consider different bones’ unique properties while studying their development and reactions to genetic changes.

In summary, this study elucidates the important role of SIKs in controlling alveolar bone osteoblast maturation. Global SIK2/SIK3 deletion results in reduced alveolar bone formation and bone mass unlike the long bone, where SIKs gene ablation leads to increased bone formation and trabecular bone mass. These alveolar bone defects are present during development, adult remodeling, and in tooth extraction models. Comparative single cell RNA-sequencing analysis indicates that alveolar bone osteoblasts exhibit distinct transcriptomic profiles compared to long bone osteoblasts. Understanding the molecular mechanisms by which SIK2/SIK3 selectively regulate osteoblast maturation in alveolar bone could have significant clinical implications. The application of small molecule SIK modulators may enhance alveolar bone quality following tooth extraction or aid in maintaining alveolar bone health in the aging population.

## Materials and methods

### Genetically modified mice

All animals were housed in the Center of Comparative Medicine at the Massachusetts General Hospital, and the Massachusetts General Hospital Subcommittee on Research Animal Care approved all of the experiments conducted on the animals. The following published genetically modified strains were used: *Sik2* floxed mice (RRID: MGI: 5905012), *Sik3*^tm1a(EUCOMM)Hmgu^ mice (RRID: MGI 5085429) were purchased from EUCOMM and bred to PGK1-FLPo mice (JAX #011065) to generate mice bearing a loxP-flanked SIK3 allele^27^. Generation of *Sik2/Sik3* floxed mice was previously described^62^; the two genes are linked on mouse chromosome 9. *Six2*-Cre mice were from Jordan Kreidberg (Boston Children’s Hospital, Boston, Massachusetts, USA). *Ubiquitin-*Cre^ERt2^ mice^63^ (JAX #008085) were intercrossed to *Sik2/Sik3* floxed mice. *Six2*-Cre mice were intercrossed to Ai14 td Tomato^LSL^ reporter mice (JAX #007914). *Cre^ERt2^*-negative littermate controls were used for all studies to account for the potential influence of genetic background and the impact of tamoxifen on bone development and homeostasis. All mice were backcrossed to C57BL/6J for at least five generations.

Tamoxifen (Sigma-Aldrich, St. Louis, MO catalog #T5648) was dissolved in 100% ethanol at concentrations of 5 mg/mL and 20 mg/mL. Equal amounts of sunflower oil (Sigma, catalog #99021-250 ML-F) were added. The mixture was kept in a 60-degree incubator without a cap overnight to allow the ethanol to evaporate. In the developing model, one-time injection of 0.25 mg tamoxifen was done. In adult mice, injection of 2 mg of tamoxifen was administered three times, every other day for a week at 12-13 weeks of age.

### Tooth extraction model

Control (*Sik2*^f/f^*;Sik3*^f/f^) and mutant (*ubiquitin-*Cre^ERt2^; *Sik2*^f/f^*;Sik3*^f/f^) mice were utilized in the experiment. At 12 weeks of age, all mice received an intraperitoneal treatment of 2 mg of tamoxifen once every other day for a week. The first maxillary molar extraction took place 10 weeks later. During the surgical procedure, mice were pre-anesthetized with 2-4% Isoflurane and oxygen via an isoflurane vaporizer until they were in lateral recumbency. After that, the animal was placed in the surgical operating area in a nose cone that supplies adequate amounts of oxygen and isoflurane to maintain a surgical plane of anesthesia. Dental extraction forceps for rodents were used to remove the first maxillary molars on both sides while the mice were under general anesthesia. These mice were sacrificed 3-4 weeks after the surgical procedure; the maxilla were collected.

### Histology

Mandibles and maxillae were meticulously dissected and then preserved overnight in 4% Paraformaldehyde (pH 7) at 4 degrees Celsius. After that, they were decalcified in 15% EDTA for 1-14 days, depending on the animal’s age. Following decalcification, samples were cryoprotected in 30% sucrose/PBS for an additional night, followed by 1:1 ratio of 30% sucrose/PBS: optimal cutting temperature (OCT). compound solution overnight. Samples were cryosectioned at a thickness of 12 µm after being embedded in OCT compound. Standard techniques were followed to perform H&E and DAPI (4’,6-diamidino-2-phenylindole) staining on some sections.

TRAP staining was used to identify osteoclasts. Sections were incubated in acetate buffer (pH 5.0) for 30 minutes at room temperature. After adding Napthol AS-MX Phosphate and Fast Red TR Salt to the mixture, sections were incubated in the solution for 20 minutes at 60 degrees Celsius. Finally, Fast Green was used to counter stain the slides.

To evaluate alkaline phosphatase activity, sections were submerged in Tris Buffer (pH 9.4) at room temperature for 1 hour, then incubated in an alkaline phosphatase staining solution (Tris buffer (pH9.4), Fast Blue solution, Napthol ASBI Phosphate solution, and magnesium chloride) at 37 degrees Celsius for 20 minutes.

### RNAscope in situ hybridization

Mandibles of C57BL/6 mice were used in this experiment. At 10 days old, mandibles were collected and fixed overnight in PFA (pH 7) at 4 degrees Celsius. Then, the mandibles were decalcified in 15% EDTA for 1 days prior to paraffin processing. In situ hybridization was performed according to the manufacturer’s instructions with RNAscope 2.5 Brown HD Assay (322310, Advanced Cell Diagnosis). Following deparaffinization, sections (5 μm) were blocked with hydrogen peroxide at room temperature for 10 minutes. Antigen retrieval was then performed using Pepsin (R2283, Sigma-Aldrich) in a HybEZ 40-degree Celsius oven for 30 minutes. *Sik1* (526411, ACD), *Sik2* (526421, ACD) and *Sik3* (526431, ACD) probes were applied to the sections and incubated in a HybEZ oven for 2 hours. *Dabp* (310043, ACD) probes were used as a negative control and *polyA* (318631, ACD) probes were used as a positive control. The signal was amplified and detected using the Brown DAP kit. Lastly, the sections were counterstained using 50% Hematoxylin staining solution.

### Fluorochrome labeling

To evaluate bone formation *in vivo*, mice were administered Calcein (20mg/kg IP) on postnatal days 10 and 12, and euthanized at P14. Mandibles were dissected, fixed in 4% PFA overnight before being cryoprotected with 30% sucrose/PBS overnight, and 1:1 30% sucrose/PBS: OCT overnight. Uncalcified samples were embedded in OCT compound and cryosectioned at 12 µm. Counterstaining was done using DAPI staining.

To visualize the mineralization tissue, Von Kossa staining was performed. Uncalcified sections were immersed in 1% Silver Nitrate under 60-Watt UV lamp for 25 minutes, washed well with water, and then incubated in 2% sodium thiosulphate for 2 minutes. Lastly, Fast Green was used as a counterstain prior to imaging.

### Immunohistochemistry

#### Anti-periostin and anti-osteocalcin immunofluorescent staining

Sections were postfixed with 4% Paraformaldehyde for 20 minutes. In order to perform Osteocalcin and Periostin immunofluorescence staining, these sections were first permeabilized with 0.1% Triton-X/TBS for 30 minutes, then blocked with 5% BSA/TBST for another 30 minutes. Sections were then incubated with anti-Osteocalcin rabbit antibody (1:200, AB93876 Abcam), or anti-Periostin rabbit antibody (1:2000, ABT280 Sigma-Aldrich) overnight at 4 degrees Celsius. After that, sections were incubated in donkey anti-rabbit secondary antibody (1:400 dilution in TBST) conjugated with AlexaFlor 488 (A21206), or 568 (A10042) overnight at 4 degrees Celsius. Lastly, DAPI staining was done prior to imaging.

For Periostin immunofluorescent staining, these sections were permeabilized with 0.1% Triton-X/TBS for 30 minutes, blocked with 5% BSA/TBST for 30 mins, then incubated with anti-Periostin rabbit antibody (1:2000, ABT280 Sigma-Aldrich), or anti-Osteocalcin rabbit antibody (1:200, AB93876 Abcam) overnight at 4 degrees Celsius. Sections then were incubated in donkey anti-rabbit secondary antibody (1:400 dilution in TBST) conjugated with AlexaFlor 488 (A21206), or 568 (A10042) overnight at 4 degrees Celsius. Finally, sections were counterstained with DAPI (4’,6-diamidino-2-phenylindole,) prior to imaging.

### Image quantification

All analyses were conducted in a blinded manner. In the developmental model, the interradicular bone area was selected as the region of interest (ROI) to represent the alveolar bone of the mandible. For the tooth extraction model, the mesial root socket was used as the ROI. The number of TRAP-positive cells was counted across four sections per sample and quantified relative to the total bone surface at P14 and the total tissue area at P28. For Alkaline phosphatase staining, Osteocalcin, and Periostin immunostaining, the proportion of positively stained area was measured and calculated relative to the total tissue area.

### Three-dimensional micro-computed tomography analysis of mouse samples

For developmental model, a microCT machine (μCT40, Scanco Medical) was utilized to scan mandible as DICOM files. The scan were performed with a 10µm isotropic voxel size, 70kVp peak potenital, 114A intensity, 0.5 mm AL filter, and 200 ms integration time. For adult and tooth extraction model, a microCT machine (μCT100, Scanco Medical) was used to scan the mandible and maxilla as DICOM files. The scan were performed with a 12µm isotropic voxel size, 70kVp peak potenital, 114A intensity, 0.5 mm AL filter, and 500 ms integration time. These DICOM files were converted into Guys Image Processing Lab (gipl.gz)” files, then three-dimensional volumetric label maps, also known as segmentation, were generated with ITK-SNAP open software. The incisor and three molars of each jaw were labeled separately using the active contour approach, which defined the borders of the label maps of each tooth based on the gray level and intensity of the image. Three-dimensional segmentations of each sample were created wherein both jaws: incisor and three molars were labeled separately using the active contour method to define boundaries of label maps of each tooth based on image gray level and intensity. Then, these segmentations were transformed into three-dimensional surface models with the Slicer program. Fiducials were placed on all molars and the inferior border of the mandible with the Q3DC tool. The linear distance between fiducials was measured quantitatively to compare the width and length of the crown and roots, the height of eruption, and the height of the alveolar bone. Measurements and fiducials were completed following a previously published study^64^. To assess the mineralization tissue in alveolar bone, the interradicular bone of the first molars was chosen as an area of interest in the development and adult mouse model. For the tooth extraction experiment, the mesial root of the maxillary first molar sockets was selected as an area of interest. We evaluate alveolar bone volume fraction (BV/TV, %) and alveolar bone mineral density (BMD). Bone was separated from soft tissue using fixed threshold of 550 mg HA/cm^3^. The scans and analysis adhered to the guidelines for using microCT to examine bone architecture in rats^65^. The microCT analysis were done blindly, with each animal assigned to coded sample numbers.

### Single cell RNA-sequencing data analysis

(GSM5623768 from GEO:GSE185794)^34^ and P25 alveolar bone dataset (GSM5140554 from GEO:GSE168450)^35^ were analyzed using Seurat. For quality control, cells with less than 1,000 genes per cell and more than 15% mitochondrial read content were filtered out for downstream analysis. Harmony pipeline^36^ was used for sample integration, normalization, dimensionality reduction analysis, clustering, and visualization of the merged dataset. Osteoblast cell clusters were identified based on marker genes *Bglap*, *Col1a1*, *Sp7*, and *Dmp1*. Cells within each osteoblast cluster were down-sampled randomly using “sample()” to equalize cell number between the two datasets for cluster-specific differential expression gene (DEG) analysis. DEGs analysis was performed using a non-parametric Wilcoxon rank-sum test with the function “FindMarkers().” Statistically significant DEGs were determined based on the criteria of adjusted p-value < 0.05 and |average log2FC| of 1. DEGs were visualized on a volcano plot. Violin plots show normalized gene expression levels of *Bglap*, Col1a1, Sik2 and Sik3 in the down-sampled osteoblast cell clusters.

### Statistical analysis

MicroCT measurement were taken, including distance between the crown tip and the most inferior part of the mandible (tooth eruption), crown and root length, alveolar bone volume fraction, and bone mineral density. Unpaired Student’s T-test was performed to compare the means of each measurement between control and mutant mice. A statistically significant difference was defined by a p-value of less than 0.05.

## Acknowledgements

We thank Drs. Yingzi Yang, Vicki Rosen, Roland Baron, Francesca Gori, Henry Kronenberg, and all members of the Wein laboratory and the Massachusetts General Hospital Endocrine Unit for helpful discussions. MNW acknowledges funding support from the NIH (R01DK116716 and R01DE030416) and support as a Chen Institute MGH Research Scholar. WO acknowledges funding support from NIH (R01DE030416). NT acknowledges support from the Harvard School of Dental Medicine Presidential Scholarship. MicroCT scanning was performed at the MGH Center for Musculoskeletal Research (NIH P30 AR075042).

